# Cellular barcoding of protozoan pathogens reveals the within-host population dynamics of *Toxoplasma gondii* host colonization

**DOI:** 10.1101/2020.08.06.239822

**Authors:** Ceire J. Wincott, Gayathri Sritharan, Henry J. Benns, Farzana B. Liakath, Carla Gilabert-Carbajo, Monique Bunyan, Aisling R. Fairweather, Eduardo Alves, Ivan Andrew, Laurence Game, Eva M. Frickel, Calvin Tiengwe, Sarah E. Ewald, Matthew A. Child

## Abstract

Molecular barcoding techniques have emerged as powerful tools to understand microbial pathogenesis. However, barcoding strategies have not been extended to protozoan parasites, which have unique genomic structures and virulence strategies compared to viral and bacterial pathogens. Here, we present a versatile CRISPR-based method to barcode protozoa, which we successfully apply to *Toxoplasma gondii* and *Trypanosoma brucei*. The murine brain is an important transmission niche for *T. gondii*, and brain persistence is a clinically untreatable feature of infection. The blood-brain barrier is expected to physically restrict parasite colonization of this niche, resulting in a selection bottleneck. Using libraries of barcoded *T. gondii* we evaluate shifts in the population structure from acute to chronic infection of mice. Contrary to expectation, most barcodes were present in the brain one-month post-intraperitoneal infection in both inbred CBA/J and outbred Swiss mice. Although parasite cyst number and barcode diversity declined over time, barcodes that represented a minor fraction of the inoculum could become a dominant population in the brain by three months post-infection. Together, these data establish the first, robust molecular barcoding approach for protozoa and evidence that the blood-brain barrier does not represent a major bottleneck to colonization by *T. gondii*.

## Introduction

Microbial colonization of tissues within the host organism is a key feature of host-pathogen interactions, and often responsible for the pathology of the associated disease^1^. For example, colonization of the brain by the eukaryotic protozoan parasite *Toxoplasma gondii* during the chronic phase of an infection can lead to toxoplasmic encephalitis in immunocompromised hosts^2^. Similarly, colonization of the brain niche by the distantly related protozoan *Trypanosoma brucei brucei* is a major contributor to trypanosomiasis pathology^3^. In the case of *T. gondii* infection, the spatial redistribution is linked to parasite differentiation. The acute phase of infection, whether initiated via the natural, per oral route or intra peritoneal inoculation, is characterized by rapid tachyzoite replication and dissemination to nearly every tissue in the host. The chronic phase of infection is accompanied by systemic immune clearance and parasite conversion to slower growing, encysted bradyzoites that persist in skeletal muscle and the brain^4^. The ‘immune privileged’ quality of the chronic infection tissue niches like the brain coupled with the protective properties of the *T. gondii* cyst wall are essential for transmission. It follows that intraspecific competition prior to and during brain colonization is expected to have an impact on long-term success of a clone. Despite recent advances in our molecular understanding of tachyzoite-to-bradyzoite differentiation^5,6^, our comprehension of *T. gondii*’s within-host population dynamics remains limited. One critical function of tissue barriers, including the blood-brain-barrier (BBB) and the intestinal epithelial barrier, is to physically restrict pathogen access^7,8^. Therefore, it is anticipated that the BBB imposes a selection bottleneck upon the *T. gondii* population as the chronic infection is established. However, we lack tools to determine if the brain is colonized infrequently by a small number of parasites that subsequently expand within the tissue niche or if the host brain broadly permissive to colonization by *T. gondii*.

DNA-based molecular barcoding has been instrumental for understanding how the host and the infection process influences genetic complexity of pathogen populations, and colonization dynamics^9,10^. Early studies using restriction site-tagged poliovirus identified a bottleneck limiting the genetic diversity of viral quasi-species transmitted to the murine brain^11^. Furthermore, studies using Wild-type Isogenic Tagged Strains (WITS) of *Salmonella* have provided key insights into selection bottlenecks experienced during gastro-intestinal and invasive bacterial colonization^12,13^. The population structure of *Salmonella* infection was found to experience dramatic selective bottlenecks during the colonization of the gut niche, with the predominant colonizing strain also the dominant strain transmitted by super shedders^14^. The generation of WITS requires the insertion of molecular barcodes into the genome of the infectious agent^12^. These cellular barcodes then function as neutral alleles, allowing the complexity of the population to be closely monitored and mapped over the course of an infection using quantitative next-generation sequencing (NGS) approaches. Unfortunately, due to technical limitations molecular barcoding has not been amenable to study the within-host population structure of protozoan pathogens.

Here, we establish two simple, scalable CRISPR-based oligo recombineering strategies to efficiently generate libraries of molecularly barcoded *T. gondii* strains. We demonstrate the versatility of our method by successfully barcoding the bloodstream form of *T. brucei*. Taking advantage of tools to infect, monitor and re-isolate *T. gondii* from the natural murine host, we define how the acute-to-chronic transition and brain colonization by *T. gondii* shapes the population structure of this eukaryotic pathogen. In tissue culture infection and in the first days of *in vivo* infection, we confirm that barcoded parasite libraries are stable, enabling quantitative analysis of the effect of the intact host organism environment upon parasite population dynamics. We then use these libraries to investigate the influence of the host organism upon the infection population structure of *T. gondii* revealing that colonization of the murine brain niche is not restricted by a stringent bottleneck at the blood-brain barrier.

## Results

### Protozoan pathogens can be molecularly barcoded with a simple CRISPR-based strategy

Inspired by WITS12, we sought to establish a versatile system to molecularly barcode eukaryotic pathogens. Previously established CRISPR-Cas9 tools for *T. gondii*^15,16^ were adapted to mediate site-specific recombination of molecular barcodes into non-essential genes that encode positive selectable markers in *T. gondii* tachyzoites or *T. brucei* trypomastigotes (**Fig. 1A**). To increase recombination efficiency, *T. gondii* lacking the non-homologous end-joining pathway (RHΔ*ku80*) were used^17^. Parasites were co-transfected with a plasmid encoding both the Cas9 nuclease and guide RNA (gRNA) scaffold^15^, and a unique 60-nucleotide single-stranded donor template encoding the molecular barcode. Successful targeting of Cas9 and genomic integration of the barcode disrupted the *UPRT* coding sequence, conferring resistance to the prodrug 5-fluorodeoxyuridine (FUDR)^18^. To prevent further modification of the *UPRT* locus following a single barcode integration event our strategy also deleted both the protospacer DNA sequence recognized by the CRISPR gRNA and the protospacer adjacent motif (PAM). We confirmed barcode integration and the expected genomic by Sanger sequencing of the *UPRT* locus in the drug-resistant parasite population (**Fig. 1B**). A similar molecular barcoding strategy was applied to the bloodstream form *T. brucei* where Cas9 was targeted to the *AAT6* locus^19^, a single-copy non-essential gene that confers sensitivity to eflornithine^20^ (**Fig. 1A**). Parasites stably expressing Cas9 were co-transfected with *AAT6*-targeting sgDNAs and a double-stranded barcoding oligo. Successful barcode integration into the *AAT6* locus conferred resistance to eflornithine. Sanger sequencing confirmed the expected genomic rearrangement and barcoding of the *T. brucei* genome (**Fig. 1B**). These data indicate our barcoding strategy is an efficient tool to insert unique a barcode sequence at specific genomic positions in two divergent eukaryotic pathogens.

**Figure 1:**
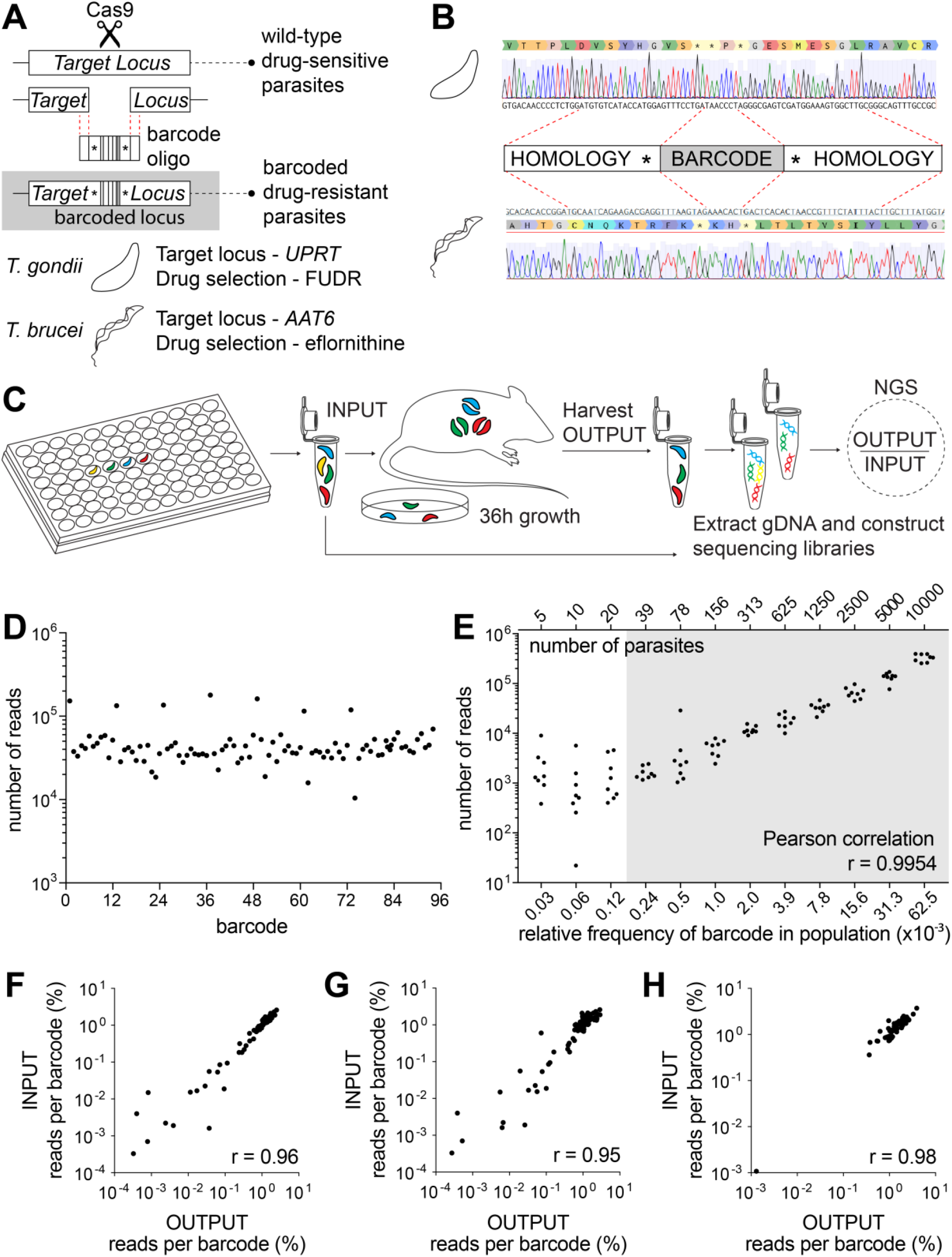
Protozoan pathogens can be molecularly barcoded using a simple CRISPR-based strategy. **A**) Schematic of molecular barcoding strategy. A CRISPR guide RNA targets Cas9 to the *UPRT* locus in *T. gondii* or the *AAT6* locus in *T. b. brucei*, where Cas9 endonuclease activity introduces a double-strand break. This break is repaired by homologous DNA repair using the co-transfected 60mer barcode oligo donor template. **B**) Sanger sequencing confirmation that barcode integration results in disruption of the *UPRT* in *T. gondii* or *AAT6* in *T. b. brucei* coding sequence and destruction of the protospacer DNA sequence and PAM. **C**) Multiplexed transfection strategy and NGS pipeline to generate barcoded libraries of *T. gondii* tachyzoites for *in vitro* and *in vivo* studies. **D**) Complex libraries containing 96 uniquely barcoded strains can be produced and the relative frequency of each barcode quantified by NGS. Scatter plot presents read counts for individual barcodes. **E**) Individual barcodes can be detected at low frequencies, and read depth is a sensitive proxy for parasite number. Scatter plot presents the number of reads for a serial two-fold dilution series of known numbers of parasites and the relative frequency of individual barcodes in the population. The shaded area indicates the data used to calculate the PCC provided: r = 0.9954, n = 9, P (two-tailed) = <0.0001. **F-G**) The population structure of multiplexed barcode libraries is stable *in vitro*. Scatter plot comparing 96 individual barcode frequencies within the INPUT pooled library population and the OUTPUT sample harvested after one ~36-hour lytic growth cycle (**F**, PCC r = 0.96, n = 96, P (two-tailed) = <0.0001), or six lytic growth cycles (**G**, PCC r = 0.95, n = 96, P (two-tailed) = <0.0001). **H**) The population structure of multiplexed barcode libraries is stable during the early stage of *in vivo* infection. Scatter plot comparing 63 individual barcode frequencies within a pooled library population for an intraperitoneal inoculum of 2×10^6^ (INPUT) and the OUTPUT sample harvested from the peritoneal cavity of C57BL6 mice 36 hours post-infection. PCC r = 0.98, n = 63, P (two-tailed) = <0.0001.

### Barcode alleles can be identified and quantified in complex populations

To examine the stability of individual barcodes across a population of uniquely labeled parasites and to evaluate the sensitivity of barcode detection, we focused on *T. gondii*. First, we generated a plate-mapped library of 96 uniquely barcoded parasites lines using multiplexed transfection (**Fig. 1C**). We reasoned that this multiplexed strategy would be useful for future chemical or genetic screening applications applied to each of the 96 uniquely barcoded strains prior to pooling. To quantify the relative representation of individual barcodes after pooling an NGS pipeline was developed. DNA was extracted from the pooled parasite library, the barcoded region of the UPRT locus was amplified for Illumina sequencing, and barcodes representation quantified using Galaxy (**Fig. 1C** and **Fig. S1A**)^21,22^. Our pipeline successfully identified all 96 uniquely barcoded strains from the pooled population (**Fig. 1D**). Notably, wells from one row of the 96 well plate were consistently more highly represented in the pooled population, reflecting a technical challenge of the 96-well transfection method (**Fig. 1D**). While unplanned, this observation suggested that the NGS readout was highly sensitive to differences in barcode frequency in the pooled population. This was confirmed experimentally, as greater variation in barcode representation was observed between biological replicate transfections (**Fig. S1B-C**) than between technical replicates of DNA indexed for NGS sequencing (**Fig. S1B and D-E**), underscoring the reproducibility of the NGS pipeline.

To determine the sensitivity of the NGS readout we generated a two-fold dilution series of the pooled parasite population prior to isolating DNA for NGS sequencing. A positive correlation was observed between the number of reads (read output) and the number of parasites in the input from 10,000 parasites/barcode down to 39 parasites/barcode (**Fig. 1E**, r = 0.9954). This confirmed that barcodes could be successfully identified and reliably quantified within libraries of at least 96-member complexity at relative frequencies as low as 0.0002 (0.02%).

### Multiplex barcoded parasite libraries are stably maintained in vitro and in vivo

To test whether the complexity of the barcode libraries was stably maintained *in vitro* the pooled library of barcoded parasites was serially passed through human foreskin fibroblasts (HFFs). Lysed-out parasite cultures were sampled every passage (~36 hours) for a period of six passages, equivalent to six lytic growth cycles (invasion, replication, egress). Relative to the input, the genetic complexity of the barcode population *in vitro* was remarkably stable after one lytic cycle or six passages (**Fig. 1F-G**). We next tested our ability to propagate and recover the barcode library *in vivo* within a murine host using a pooled library of 63 barcodes (in this pilot experiment some wells were not efficiently transfected). An inoculum of 2× 10^6^ parasites was injected intraperitoneally into C57BL/6 mice. After 36 hours parasites were isolated from the peritoneal cavity by lavage and compared to the input population. A strong positive correlation was observed between the input and peritoneal exudate populations (**Fig. 1H**, PCC r = 0.98), demonstrating that the genetic complexity of the multiplex barcode library was stable over the first 36 hours of *in vivo* infection.

### Libraries of barcode alleles can be generated by a ‘one-pot’ transfection method

One challenge of the multiplex platform arose from experiment-to-experiment variability in transfection efficiency across individual wells (**Fig. 1D and 1H**). Our CRISPR-Cas9 strategy was designed to ensure the integration of a single barcode into the UPRT locus by simultaneously deleting the protospacer and PAM motifs recognized by the CRISPR gRNA. We hypothesized that a barcoded library of parasites could be generated by a single transfection with the Cas9-gRNA plasmid and a pooled library of oligonucleotide repair templates using a widely available cuvette-based transfection apparatus (**Fig. 2A**). This ‘one-pot’ method was tested on type I RHΔ*ku80* and type II PruΔ*ku80* parasite strains in parallel, using the same mixed pool of 96 barcode oligo repair templates. FUDR-resistant parasite populations were enriched, genomic DNA isolated, UPRT locus amplicons prepared, and NGS libraries sequenced. All 96 barcodes were identified in the sequence reads generated from the UPRT locus amplicon (**Fig. 2B**). When the distribution of barcodes in RHΔ*ku80* and PruΔ*ku80* were compared directly, the two independently transfected parasite strains exhibited correlated frequency distributions for all but the least abundant barcodes, with a Pearson correlation coefficient of 0.7 (**Fig. 2B**). Similar to the multiplex method we confirmed the technical reproducibility of NGS runs by comparing technical replicates of indexed DNA amplicons (**Fig. S2**). We concluded that differences in barcode frequency most likely reflect minor variations in the relative abundance of each barcode oligonucleotide template within the pool used for transfection, rather than a negative impact of some barcodes on parasite fitness. It is notable that a higher level of biological reproducibility was observed for one-pot library transfections than the original multiplexed transfection approach (**Fig. S1C**). We next tested the one-pot strategy in *T. brucei*. We were similarly able to produce complex 96-member libraries of uniquely barcoded strains from a single transfection (**Fig. 2C**). The one-pot transfection protocol was used for all subsequent experiments.

**Figure 2:**
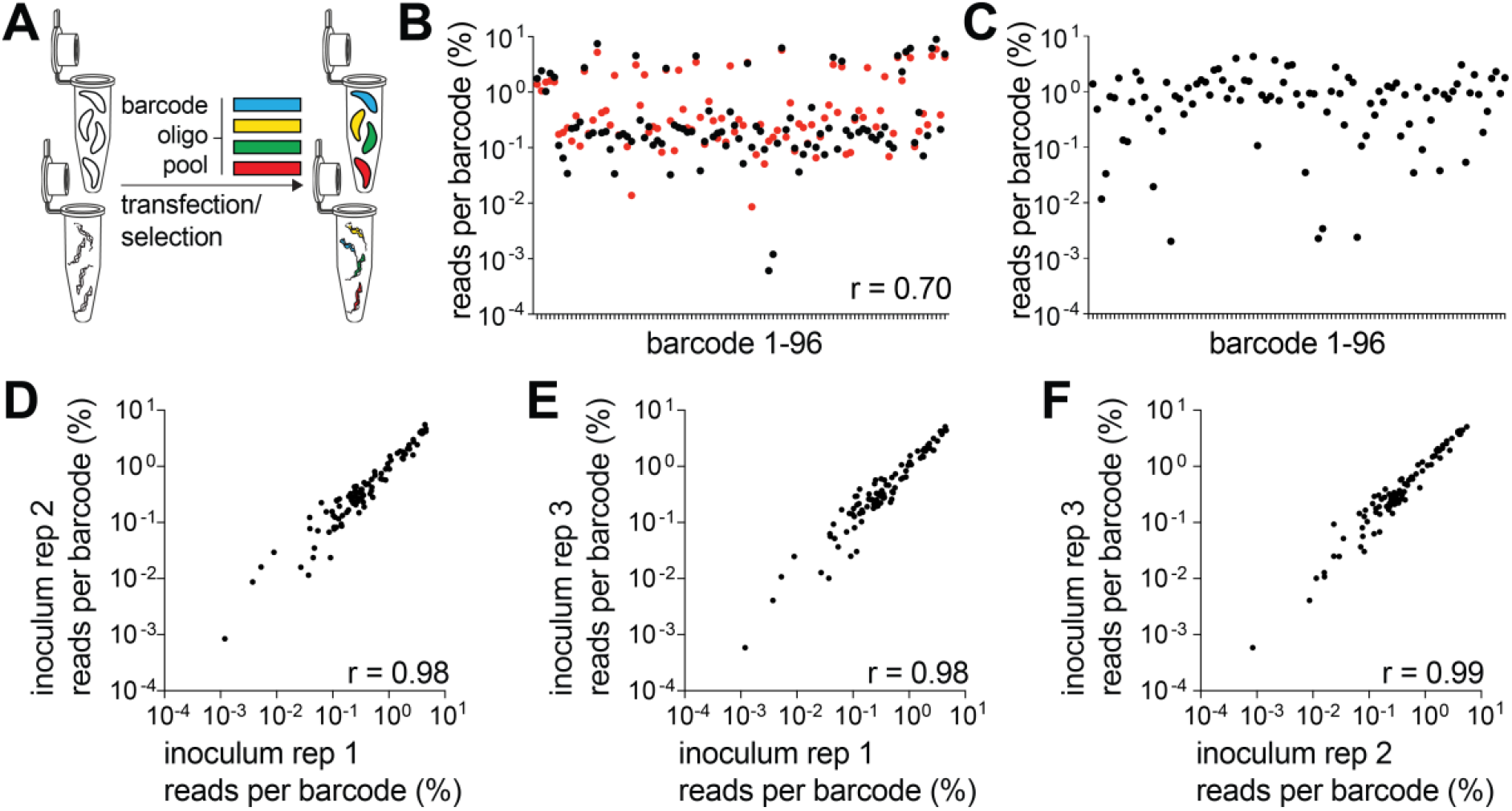
Barcoded libraries can be generated via one-pot transfections. **A**) Schematic of the one-pot transfection method to generate barcoded parasite libraries. **B**) Overlay of the relative percent frequency of each barcode in an RHΔ*ku80* population (black dots) or a PruΔ*ku80* population (red dots) generated from a common pool of barcode oligos by one-pot transfection. The scatter plot is distributed according to barcode identifier (1-96) and PCC compares the two populations, r = 0.70, n = 96, P (two-tailed) = <0.0001. **C**) Data from one-pot transfections with *T. brucei*. Scatter plot presents relative percentage frequency of barcodes within transfected parasites, distributed according to barcode identifier (1-96). **D-F**) One-pot transfections using PruΔ*ku80* were diluted to a founder population of 37,000 parasites and re-expanded in HFFs prior to NGS. All 96 barcodes were identified in each sample and the relative percent frequency of barcodes was highly correlated in pairwise comparison each inoculum sample: **D**) inoculum 1 vs. 2, PCC r = 0.98, n = 96, P (two-tailed) = <0.0001; **E**) inoculum 1 vs. 3, PCC r = 0.98, n = 96, P (two-tailed) = <0.0001; **F**) inoculum 2 vs. 3, PCC r = 0.99, n = 96, P (two-tailed) = <0.0001. PCC values provided on scatter plots indicate degree of correlation between populations being compared.

### *Cellular barcodes reveal the population structure of a* T. gondii *infection in vivo*

The transition from acute to chronic infection corresponds with the spatial redistribution of *T. gondii* to skeletal muscle and the central nervous system^4^. This is accompanied by tachyzoite differentiation into bradyzoites cysts, which are necessary for parasite transmission^5^. The restrictive nature of the BBB is well documented in other infection models^23^. We hypothesized that these spatial, temporal, and developmental transitions would impose bottlenecks upon the parasite population represented in the brain at chronic infection. We sought to test this by infecting murine hosts intraperitoneally with the pooled library of PruΔ*ku80* parasites stably expressing 96 neutral barcode alleles.

First, we confirmed that diluting the PruΔ*ku80* library to a dose tolerated by mice did not disrupt the population structure in a way that could confound the interpretation of *in vivo* experiments. Three inoculum samples of 37,000 viable tachyzoites (determined by plaque assay) were plated on HFFs and re-expanded in tissue culture followed by DNA isolation for NGS sequencing. In each inoculum sample, all 96 barcodes were detected by NGS sequencing and pairwise comparisons of each sample were strongly correlated (**Fig. 2D-F**), confirming that *in vitro* expansion of *in vivo* samples would minimally affect library composition.

An *in vivo* experiment was conceived to study the effect of murine host colonization. This would probe the within-host population genetics of the *T. gondii* infection as it progressed from the initial acute infection in the peritoneum to the chronic infection in the brain (**Fig. 3A**). To determine if the parasite population was stable early in acute infection, parasites were isolated from the peritoneal cavity of some mice at 48 hours post-infection and expanded in vitro. Although barcode extinction was rare (one barcode in a single mouse), the relative frequency of barcodes in each peritoneal isolate was unique to each animal (**Fig. 3B-C**). This observation supports a model where the initial selective sweep at the onset of acute infection is stochastic and emphasized the need to consider each host organism as a unique environment. After 28 days, the brains of chronically infected mice were isolated, and parasites expanded *in vitro*. Unexpectedly, the majority of barcodes were detected in each host brain (**Fig. 3D, Table S1**), consistent with a minimal founder effect on the genetic diversity of the parasite population colonizing the central nervous system. The cumulative extinction frequency across all 14 mice was 0.007 (10 of 1,344 total barcodes) and lost barcodes were most often represented at low abundance within the inoculum (**Table S3**). Of note, these extinction events were observed in the NGS runs with the lowest total read counts, suggesting that extinctions could be due to reduced sequence sampling depth rather than a true absence of the specific barcoded parasite(s) in the brain (**Table S3**). Together, these data indicate that any selection bottleneck experienced by parasites during the colonization of the brain niche must be broad.

**Figure 3:**
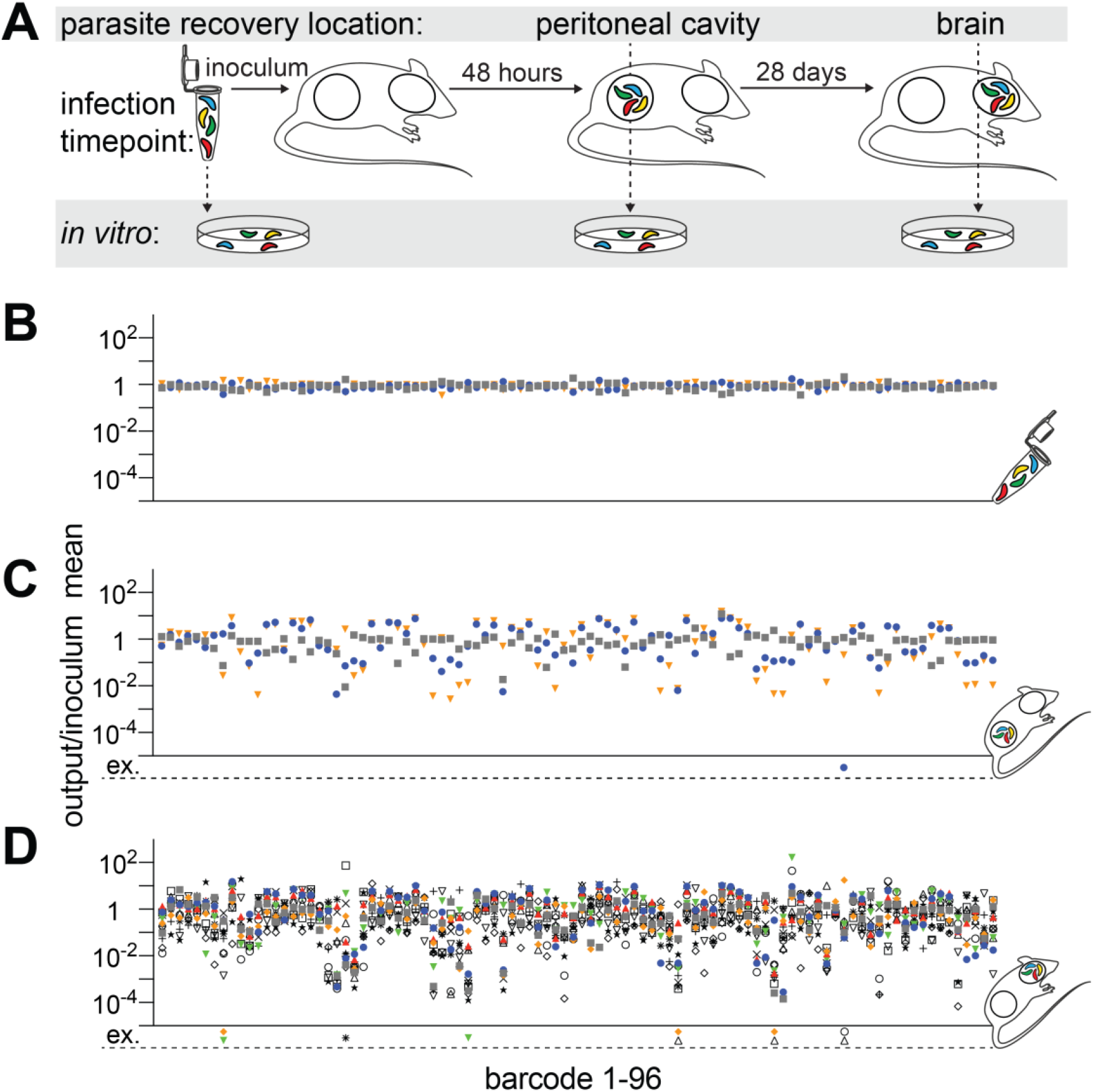
The murine host brain is permissively colonized by *T. gondii*. **A**) Schematic of an experiment to identify changes in the *T. gondii* population structure over the course of a 28-day *in vivo* infection. CBA/J mice were infected with an inoculum of 37,000 tachyzoites injected intraperitoneally. The infectious population structure was monitored in peritoneal exudates at 48 hours post-infection (early acute) or in the brain at the onset of the chronic phase (28 days post-infection). Parasites were re-expanded in tissue culture prior to NGS sequencing. **B-D**) Scatter plot represent individual barcode frequencies in each sample relative to the mean of the inoculum in the inoculum replicate (**B**, N=3 replicates), the peritoneal cavity at 48 hours (**C**, N=3 mice), and the brain at 28 days (**D**, N=14 mice). Barcode extinctions (ex.) are indicated below the x-axis in the corresponding position to the absent barcoded strain (see also Table S1). Each symbol represents an individual inoculum or mouse.

#### Stochastic selection and host genotype shape the T. gondii population structure colonizing the brain

We next sought to determine how host genetic background influences the dynamics of parasite brain colonization over time. To do this we infected Swiss Webster mice, which are outbred to maximize genetic diversity and heterozygosity, and inbred CBA/J mice (**Fig. 4A**)^24^. Mean parasite cyst burden was not significantly different between CBA/J mice and Swiss Webster mice at one-month post-infection, indicating that a similar number of parasites were accessing the brain niche (**Fig. 4B**). As expected, cyst burden declined over time, but the reduction in mean cyst number was similar between mouse genotypes^24,25^.

**Figure 4:**
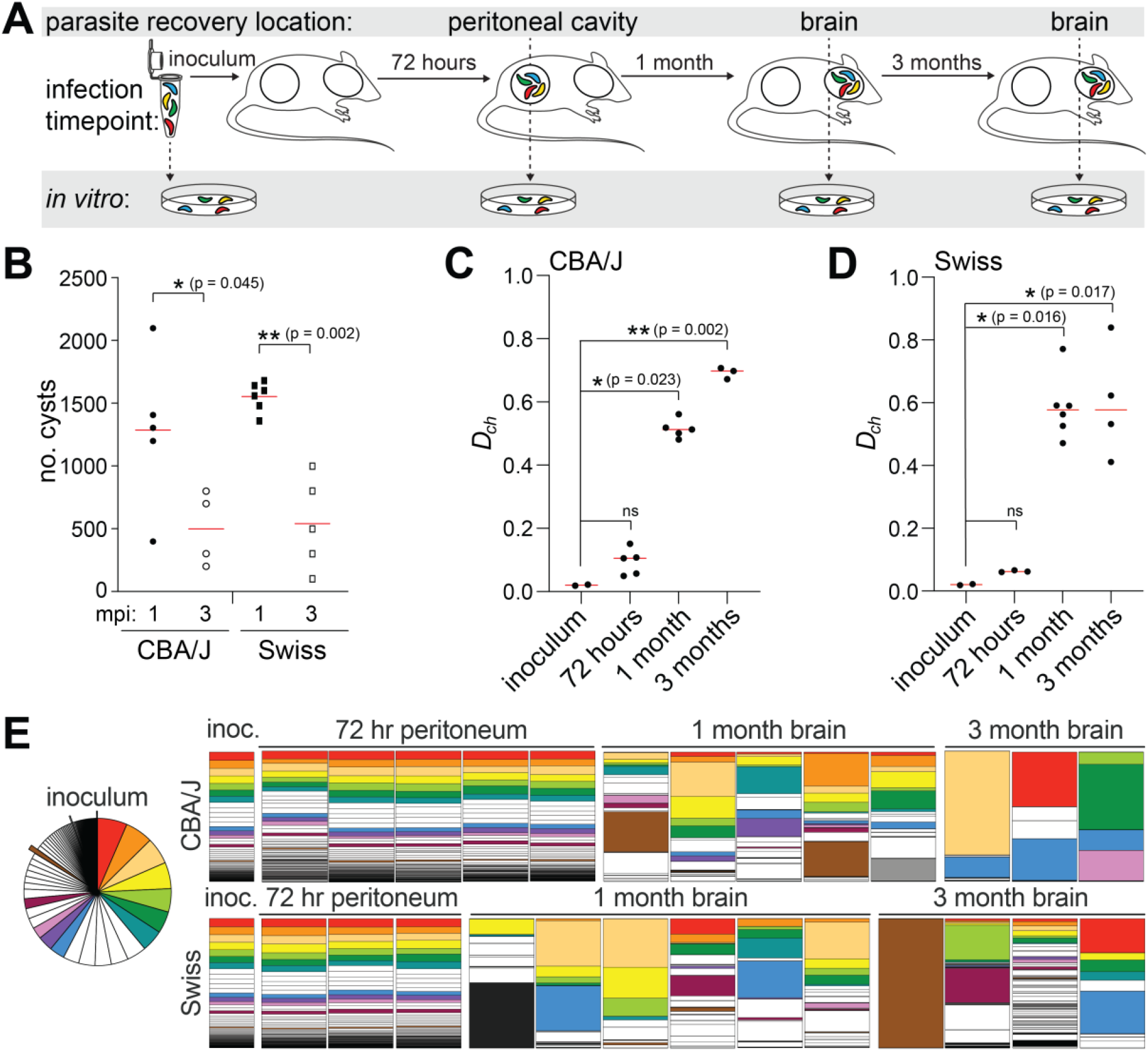
The population structure of the *T. gondii* chronic infection is dynamic within the murine host brain niche. **A**) Schematic of an experiment investigating changes in the *T. gondii* population structure over the course of a three-month *in vivo* infection in inbred CBA/J and outbred Swiss Webster mice. Mice were intraperitoneally injected with 12,000 tachyzoites and the *T. gondii* population structure was evaluated at an early acute (72 hours in peritoneal exudate), early chronic (one month in the brain), and late chronic (three months in the brain) infection time points. **B**) Bradyzoite cyst counts from *T. gondii* infected brains isolated from CBA/J and Swiss Webster mice at one and three months post-infection (mpi). Significance was tested with one-way ANOVA. **C-D**) Scatter plot of chord distances (*D_ch_*) calculated from barcode representation in the inoculum relative to 48-hour peritoneal samples, 1-month post infection brains or 3-month post infection brains in CBA/J (**C**) or Swiss Webster mice (**D**). Significance relative to inoculum was tested by Kruskal-Wallis one-way ANOVA, with Mann-Whitney used for pairwise comparisons. **E**) Parts-of-whole charts representing the relative frequency of each barcode within the mouse samples. Barcodes are ranked in descending order of abundance in the inoculum mean (inoculum mean N=3, represented as a pie chart and bar charts labeled ‘inoc.’). A colour was assigned to a barcode if it was the dominant barcode in any brain sample or if it represented greater than 20% of all reads in any brain sample. Each colour is unique to one barcode. Any barcode that dominated a mouse brain and was represented in the bottom 25% of inoculum reads is outset in the inoculum pie chart for clarity and shaded brown, grey, or black in parts-of-whole charts.

To evaluate the parasite population structure within these hosts we used Cavalli Sforza’s chord distance calculation^26^. Chord distance is a geometric measure of population distance. While chord distance does not provide absolute quantification of the width of any bottleneck, it can be used to infer changes in the genetic structure of a population between points A and B. In CBA/J mice (**Fig. 4C**) there was minimal distance separating the inoculum from the peritoneal exudate population 72 hours post-infection. A significant increase in distance was observed between the inoculum and the one-month brain samples. Consistent with the decline in cyst number over time, chord distance was greater at three months post-infection in CBA/J brain. Outbred Swiss Webster mice also exhibited little shift in chord distance from the inoculum to the peritoneal exudate and a significant shift in distance from the inoculum to the one-month post-infection brain samples (**Fig. 4D**). However, the individual Swiss mice exhibited greater variation in parasite population structure at each chronic time point than CBA/J mice. Unlike CBA/J mice, Swiss mice supported parasite populations with a similar median chord distance between inoculum and one or three months post-infection brains.

Cumulatively, our data indicated that following an IP infection the murine brain was unexpectedly permissive to colonization (**Fig. 3D**) and parasite population structure was dynamic within individual hosts (**Fig. 4C-D**). In CBA/J mice, the chord distance was greater at three months post-infection than at one-month post-infection (**Fig. 4C**), so we became interested in understanding if the barcoded lineages that were the most abundant in the inoculum had a competitive advantage for long-term persistence. In CBA/J mice, dominant barcode alleles at three months post-infection (**Fig. 4E**, top row) corresponded to parasite populations that were among the top 35% of reads (top six most abundant barcodes) in the inoculum. While this trend was similar at the one-month post-infection time point, several CBA/J mice harboured a dominant barcode lineage that corresponded to one represented in the bottom 25% of reads in the inoculum (**Fig. 4E** outset in the inoculum pie chart). In agreement with the chord distance analysis results, individual Swiss mice had diverse dominant population structures at three months post-infection (**Fig. 4E**, bottom row). Dominant barcodes in the Swiss brains were often but not always derived from lineages represented within the top 75% of reads in the inoculum (**Fig. 4E** most abundant 15 barcodes), however, this was not always the case. For example, one mouse was dominated by a barcode that was present at low frequency in the inoculum, while another mouse exhibited a population structure where barcode frequency was more evenly distributed than in the inoculum or peritoneal parasite isolates. This heterogeneity was also reflected in the barcodes that dominated Swiss brains at one-month post-infection, suggesting that even relatively low-frequency members of a founder population can establish and maintain persistent infection in a manner that likely depends on stochastic variables and host genotype.

## Discussion

Brain residency has long been appreciated as a strategy for pathogenesis exploited by *T. gondii* to evade sterilizing immunity, as well as an evolutionary strategy to increases the likelihood of transmission via predation. Feline consumption of high-fat, energy rich infected neuronal tissue of prey, including mice, provides *T. gondii* with a route back into the definitive feline host^27,28^. A means to evade any restrictive bottleneck when colonizing the brain niche would be consistent with this evolutionary strategy, supporting maximal transmission of genetic diversity into the feline host to contribute to recombination in the subsequent sexual cycle. Additionally, *T. gondii* can infect most warm-blooded animals, indicating that intermediate host-specific selection environments will frequently and unpredictably change, favouring different phenotypes at different times. It therefore likely benefits the parasite to maintain genetic diversity and phenotypic plasticity to ensure survival in varied future host species.

A key role of tissue barriers is to restrict pathogen access to the underlying tissue niche^7^. Unexpectedly our findings imply multiple, unique *T. gondii* colonization events from circulation into the brain parenchyma, rather than rare brain invasions coupled with expansion within that niche. During intraperitoneal infection, the loss of barcodes is stochastic, and most barcodes are identified in the brain parenchyma at one month post-infection (**Fig. 3**). Chord distance indicated a change in the genetic structure of the parasite population in the CNS at one month post-infection (**Fig. 4C-D**). In CBA/J mice and Swiss mice, the dominant barcodes in the brain following intraperitoneal infection tended to be those most frequently observed in the inoculum, however, most barcodes were still detected indicating limited selective pressure imparted by the BBB. However, there were exceptions to this rule, particularly in outbred Swiss mice, where minor constituents of the inoculum were able to dominate the brain population in some individuals.

We employed intraperitoneal infection because it facilitates a precise quantification of barcode frequency and coverage across the inoculum dose. This is not possible using bradyzoite cysts as each cyst, which must be harvested from infected mice, has a variable parasite number. In the future, it will be interesting to investigate how host colonization following oral infection shapes the population structure of the parasite, and how it differs from the intraperitoneal route. It should also be noted that the results from our study do not mean that there is *no* bottleneck for infection of the brain niche, simply that with the number of barcode markers used in this study we were not able to determine the bottleneck width.

The observations made in this study would have not been possible without a means to barcode this eukaryotic pathogen. Molecular barcoding has provided critical insights into the infection biology of viruses such as poliovirus^11,29^, bacteria such as *Salmonella*^12–14,30^, and *Escherichia coli*^31^. We anticipate that barcoded *T. gondii* strains will have similar far-reaching applications. While barcode sequencing strategies have been leveraged for eukaryotic pathogen phenotypic screens^32–34^, to our knowledge this is the first use of cellular barcodes to study the within-host infectious population structure of a eukaryotic pathogen. The versatility of the strategies presented will allow researchers to work with individual barcoded strains in isolation prior to pooling (*via* plate-based library generation), or with complex libraries of barcoded strains generated through our one-pot approach. We believe this will expand the application of our approach beyond the population genetic studies described herein. It should be noted that disruption of the *UPRT* locus has been documented to negatively impact cyst burden, which would suggest that our data may *underrepresent* barcode diversity in the CNS. However, all barcodes are inserted at the same position, so the cross-comparison of populations is internally controlled in this study. Our simple oligo barcoding strategy can be applied to systems accessible to CRISPR that contains endogenous or engineered negative selection markers, which we successfully demonstrated with the first cellular barcoding of *T. brucei*. In principle, our approach allows for parallel integration of even greater numbers of unique barcodes, which will provide increased precision in future host-pathogen population genetic studies. An increased number of molecular barcodes combined with methods such as Sequence Tag Analysis of Microbial Populations (STAMP)^35^, or the recently published update STAMPR^36^, will make it possible to quantify the absolute founder population number present within the host brain. Combined with these methods, our molecular barcoding approach will allow researchers to probe the within-host population genetics of tissue colonization during a *T. gondii* infection with an unprecedented degree of molecular resolution.

## Methods

### Parasite cell culture

*T. gondii* parasite strains were maintained by serial passage in confluent human foreskin fibroblasts (HFF-1 ATCC® SCRC-1041™). HFFs were cultured at 37°C with 5% CO2 in Dulbecco’s Modified Eagle’s medium supplemented with 10% foetal bovine serum and 2 mM L-glutamine. Tachyzoites were harvested via mechanical syringe lysis of heavily infected HFFs through a 25-gauge needle. RHΔ*ku80* parasites were used for *in vivo* and *in vitro* studies. PruΔ*ku80* parasites were used in *in vivo* experiments where chronic infections were established. Parasite strains were received as a kind gift from Dr. Moritz Treeck.

### Generation of barcoded *T. gondii* strains and libraries

60-nucleotide single-stranded oligos were designed to include a unique six nucleotide barcode sequence flanked by a stop codon and homology regions on either side. Barcodes were designed using the DNA barcode designer and decoder, nxcode (http://hannonlab.cshl.edu/nxCode/nxCode/main.html). The sequences of all oligos within the 96-member library can be found in **Table S2**. Barcoded libraries of tachyzoites were generated using two alternative strategies: For strategy A, 96 independent transfections were carried out in 16 well Nucleocuvette strips. 10 μg of the pSAG1::Cas9-U6::sgUPRT vector^15^ and 10 μg of the barcode oligo (equivalent to an ~1:160 molar ratio of plasmid to oligo) were co-transfected into approximately 1×10^6^ extracellular tachyzoites using the 4D-Nucleofector X Unit programme F1-115 (Lonza). 24 hours post-transfection, transgenic barcoded parasites were selected for using 5 μM 5’-fluro-2’-deoxyuridine (FUDR). Barcoded strains were independently maintained, and only pooled just prior to use. For strategy B, a single “one-pot” transfection was carried out. An oligo library pool containing roughly equal amounts of all barcode oligos was prepared. The ratio of the pSAG1::Cas9-U6::sgUPRT vector to the total oligo pool was the same as in strategy A, though here the final concentration of any single oligo within the pool was ~100-fold less. Transfection and selection was performed as for A, with the complex barcoded strain library generated and maintained as a single population.

### Generation of barcoded *T. b. brucei* strains

60-nucleotide double-stranded oligos were designed to include a unique six nucleotide barcode sequence (bold, upper case) flanked by a stop codon (lower case) and 24 bp homology regions on either side: TGCAATCAGAAGACGAGGTTTAAGtag**AAACAC**tgaCTCACACTAACCGTTTCGATTTAC. Amino acid transporter *AAT6* (Tb927.8.5450) was selected as a suitable locus for barcode integration. DNA encoding the sgRNA sequence targeting the *AAT6* locus (GTTTAAGTTCACATTGTCGC) was generated by PCR as described in^19,37^, ethanol precipitated, and 10 μg was mixed with 10 ng pre-annealed oligonucleotides. The mixture (20 μl total volume) was added to ~10^7^ Cas9 expressing bloodstream form Lister 427 *T. b. brucei* cells in 100 μl Amaxa buffer and electroporated using Amaxa Nucleofector IIb (Lonza) program X-001. Transfected cells were immediately added to pre-warmed HMI-9 medium containing 270 μM eflornithine. Transfected cell cultures were passaged under selection every two days. Drug resistant parasites were harvested seven days after transfection for genomic DNA isolation. PCR amplicons encompassing the barcoded region of the *AAT6* locus (from four independent transfections) were generated using ORF-specific PCR primer sequences (5’ to 3’) ATGAGAGAGCCGATACAAACTTCAAC and TCAGAGTTCAGCAATGACGCTG. Barcode integration was confirmed by Sanger sequencing of these amplicons. For the one-pot transfection strategy, complementary single-stranded barcoding oligos were annealed to produce 96 unique double stranded barcoding repair templates. Annealed barcoding oligos were then pooled, and used in a single transfection as described above.

### NGS library preparation

Frozen cell pellets of extracellular tachyzoites were thawed to room temperature and genomic DNA extracted using the DNeasy Blood & Tissue Kit (Qiagen). Genomic DNA libraries were prepared following the 16S Metagenomic Sequencing Library Preparation guide (Illumina). In brief, an ~300 base pair amplicon region encompassing the 6 nt barcode sequence was amplified (30 cycles) from the barcoded *UPRT* locus using primer sequences (5’ to 3’) **TCGTCGGCAGCGTCAGATGTGTATAAGAGACAG**tggatgtgtcataccatggagtttcctg and **GTCTCGTGGGCTCGGAGATGTGTATAAGAGACAG**tgttttagtgtaacaaagtggacagcagc. These primer sequences include the specified Illumina adapter overhang sequences (bold, uppercase). AMPure XP beads were used to purify the resulting PCR product. An indexing PCR (10 cycles) was carried using the purified product as the template to add dual indices and sequencing adapters to the amplicon using the Nextera XT Index Kit (Illumina). Indexed libraries were then cleaned using AMPure XP beads and quantified on the Quantus Fluorometer using the QuantiFluor ONE dsDNA System (Promega). Amplicons were purity-checked and sized on a TapeStation using D1000 ScreenTape System (Agilent). For each NGS run, typically 8 to 25 uniquely indexed libraries were pooled at equimolar concentrations for multiplexed outputs on either an Illumina MiSeq or NextSeq sequencer using the MiSeqV3 PE 75 bp kit or NextSeq 500/550 Mid Output v2.5 PE 75 bp kit respectively. PhiX DNA spike-in of 20% was used in all NGS runs. Following acquisition, sequencing data was demultiplexed and total sample reads extracted from fastq files using the Galaxy web platform (www.usegalaxy.org). Within Galaxy, sequencing reads were concatenated, trimmed, and split into the respective barcodes. Phred QC scores for all NGS runs were >30 with the exception of a single run used for analysis of technical and biological replicates, which still gave an acceptable score of 28. Following trimming to the appropriate 6 nt region a stringent barcode mismatch tolerance of 0% was applied, typically resulting in 10-15% of total reads being discarded. Barcode read data was analysed using Prism 8 and correlations coefficients calculated within the software using Pearson analysis. Testing the sensitivity of the NGS pipeline (Fig. 1e), a 96 well plate of 96 uniquely barcoded strains was set up (using transfected drug resistant parasites), with 10,000 parasites/well. For the plate (12 columns x 8 rows), a serial two-fold dilution was performed across the 12 columns, and all rows in the final column pooled after the final dilution. Genomic DNA was then prepared from the final pooled sample, and processed for NGS as described.

### Mice

Six-week old female C57BL/6 or CBA/J mice were purchased from Jackson Laboratories. Mice were acclimated for seven days prior to infection. Six to eight-week old female Swiss Webster mice (originally form Jackson Laboratories) were obtained from the University of Virginia Center for Comparative Medicine foster and sentinel colony. For studies using CBA/J and Swiss Webster mice, the animal protocols were approved by the University of Virginia Institutional Animal Care and Use Committee (protocol # 4107-12-18). All animals were housed and treated in accordance with AAALAC and IACUC guidelines at the University of Virginia Veterinary Centre for Comparative Medicine. The procedures involving C57BL/6 (multiplex experiment) mice were approved by the local ethical committee of the Francis Crick Institute Ltd, Mill Hill Laboratory and are part of a project license approved by the Home Office, UK, under the Animals (Scientific Procedures) Act 1986.

### Mouse infections

For intraperitoneal infection the pooled barcode parasite library was expanded on HFFs in a T175 flask. Once full parasite vacuoles were observed, parasites were scraped and syringe lysed, counted on a haemocytometer and diluted to an inoculum of 37,000 viable parasites (data represented in Figure 3) or 12,000 viable parasites (data represented in Figure 4) in 200 μL of PBS per mouse. The numbers of viable parasites in the IP infection inoculums were determined by plaque assay. At the time of inoculation 2×10^6^ parasites were frozen as an initial population control. In addition, three inoculum control samples were expanded immediately on HFF T25 flasks. After 48 or 72 hours three to five mice were euthanized to isolate parasites in the peritoneal exudate. Specifically, 10 mL of PBS was injected by 25G needle into the peritoneal cavity, mice were rocked vigorously, and peritoneal fluid removed by syringe. Parasites and exudate cells were washed twice in 10 mL of media containing penicillin/streptomycin, pelleted at 1,500 rpm and plated on HFFs T25 flasks. Parasites were harvested when they approached full lysis of the monolayer pelleted and frozen for genomic DNA isolation. After 28 days or three months, the remaining mice were euthanized. Carcasses were incubated in 20% bleach for 10 minutes and the brain was excised in the biosafety cabinet under sterile conditions. To isolate parasites the brains were mashed though a 70 μm filter using 25 mL PBS with 5% FBS and penicillin/streptomycin. Brain mash was pelleted for 10 minutes at 1,500 rpm, washed twice with PBS and penicillin/streptomycin then plated on HFF monolayers in T75 flasks. After 36 hours, media was changed to remove debris. Parasites were harvested by syringe lysis when the HFF monolayer was nearly lysed out (approximately two weeks), pelleted and frozen for genomic DNA isolation. To confirm cyst formation in the brain at one month (28 days) or three months post infection, 1/50th of the mash was reserved, fixed in 4% paraformaldehyde for 15 minutes then stained with a 1:500 dilution of dolichos bifluorus agglutinin conjugated to FITC in PBS (Vector Labs). FITC-positive cysts were confirmed by fluorescence and morphology under 20x magnification and the total cyst burden per brain was back-calculated.

### Bottleneck Analysis and Chord Distance Calculations

Genetic selection bottlenecks experienced within the murine host were estimated by calculating changes in the relative frequencies of barcodes within dynamic *T. gondii* populations in relation to the starting population in the inoculum. The following equations 26 were used to calculate chord distance:

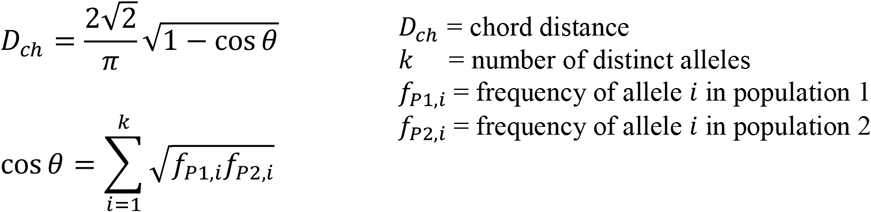

## Acknowledgements

This work was supported by grants NC/S001239/1 from the NC3Rs (to C.J.W and M.A.C), 202553/Z/16/Z from the Wellcome Trust & Royal Society (to M.A.C.) and NIH NIGMS R35GM138381 (to S.E.E). CGC and CT were supported by Wellcome Trust & Royal Society award 208780/Z/17/Z. This work was also supported by the Francis Crick Institute (FC001076 to EMF), which receives its core funding from Cancer Research UK, the UK Medical Research Council, and the Wellcome Trust. EMF was supported by a Senior Wellcome Trust Fellowship (217202/Z/19/Z). The Imperial BRC Genomics Facility provided resources and support that contributed to the research results reported within this paper. The Imperial BRC Genomics Facility is supported by NIHR funding to the Imperial Biomedical Research Centre. We would also like to thank Dr Ellen M. McDonagh for critical reading of the manuscript.

## Author Contributions

Conception/design of the work: EMF, CT, SEE, MAC. Acquisition/analysis/interpretation of data: CJW, GS, CGC, HJB, FBL, CGC, MB, ARF, EA, IA, LG, SEE, MAC. Manuscript drafting and revision: CJW, GS, HJB, EA, EMF, CT, SEE, MAC.

## Supplemental Material

**Table S1:**
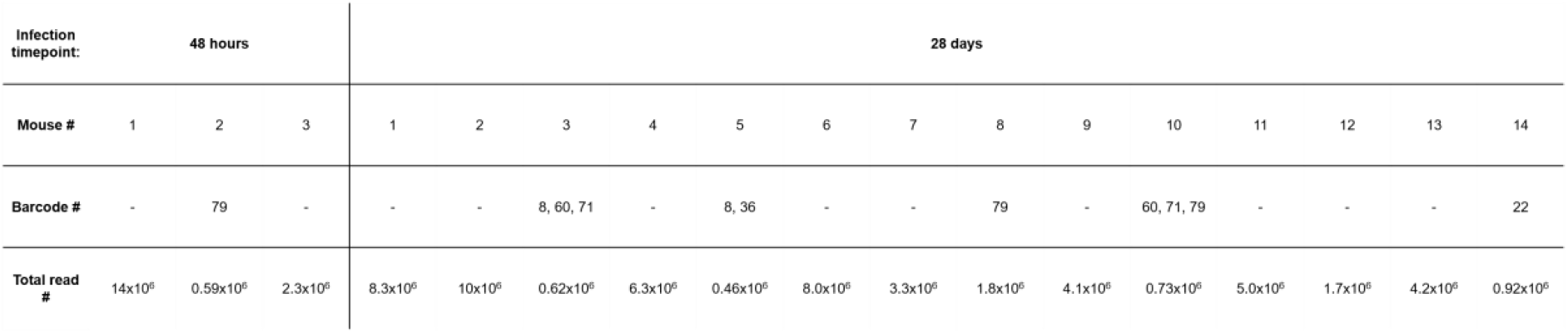
Details of barcode extinctions occurring during the different phases of infection. Barcode numbers and the mouse in which the extinction was observed are noted as shown in Figure 2C-D. Extinctions are defined by an absence of the barcode sequence within the processed NGS read data.

**Table S2:** Sequences of barcoding oligo nucleotides.

**Table S3:** Raw data from all NGS experiments.

**Figure S1:**
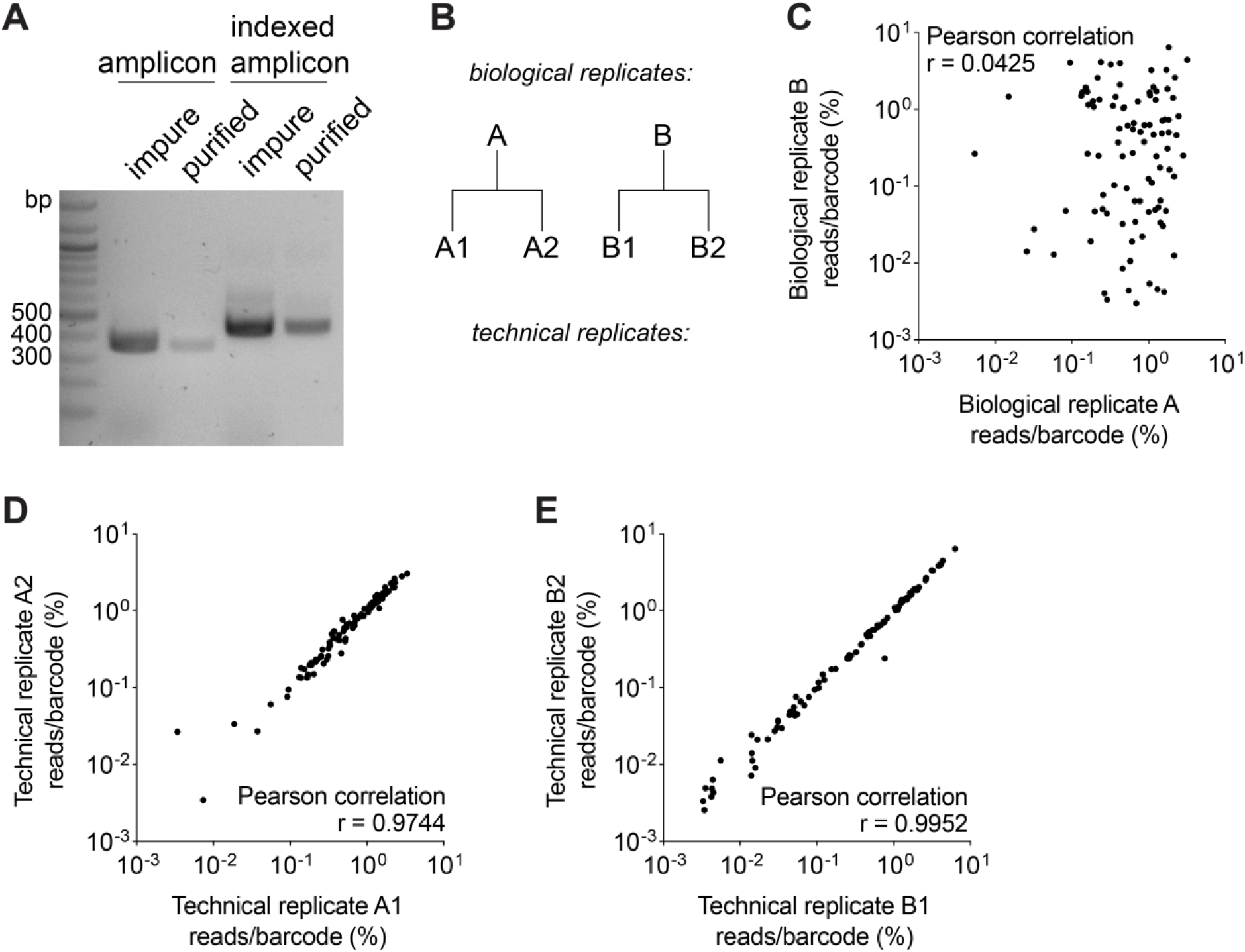
A robust pipeline for analysis of barcoded *T. gondii* libraries by NGS. **A**) A single ~300 bp region of the UPRT locus was amplified from genomic DNA and purified (amplicon, lanes 1 and 2). The purified amplicon was then indexed and re-purified prior to quantification and sizing (indexed amplicon, lanes 3 and 4). **B**) Strategy to isolate the primary source of variation to the multiplexed transfection barcoding strategy or the NGS pipeline. **C-E**) Scatter plots show the percent representation of individual barcodes within library pools for biological replicate transfections, (**C**, PCC r = 0.0425, n = 96, P (two-tailed) = 0.6807), or technical replicates of amplicon indexing (**D**, PCC r = 0.9744, n = 96, P (two-tailed) = <0.0001) and (**E**, PCC r = 0.9952, n = 96, P (two-tailed) = <0.0001). PCC values represent comparison between samples indicated on each x and y axis.

**Figure S2:**
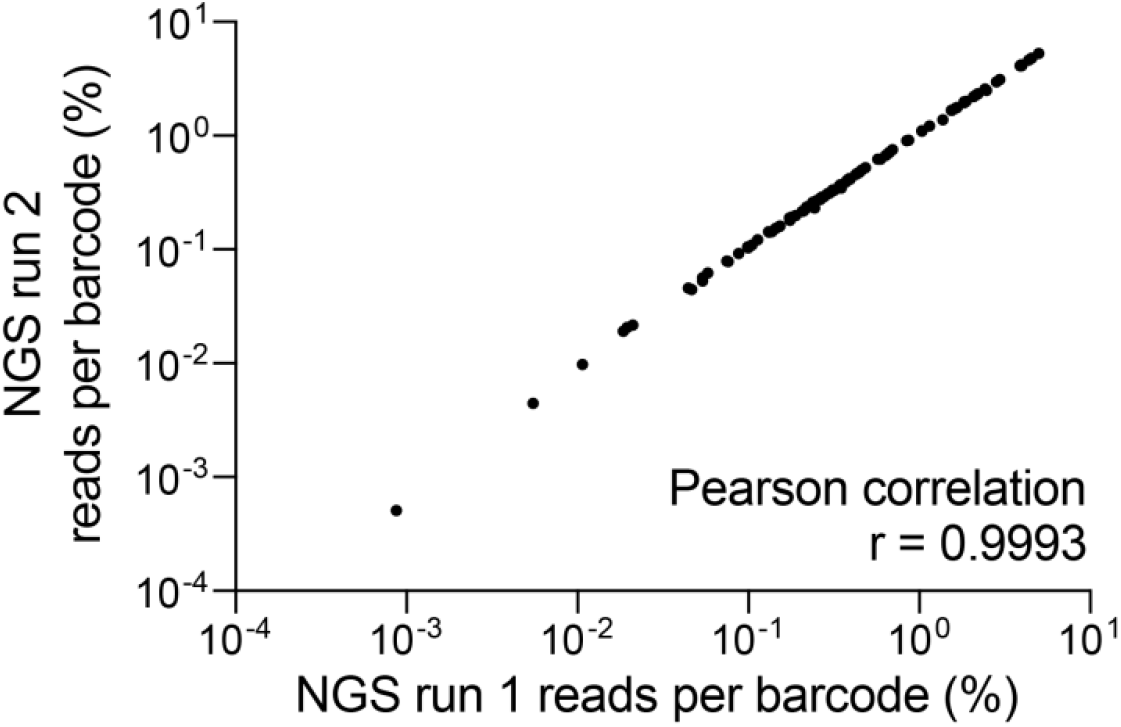
Independent NGS runs assessing barcoded strain frequency distributions are highly reproducible. Scatter plot of relative percentage frequency of barcodes within a single genomic sample processed on independent NGS runs. PCC r = 0.99, n = 96, P (two-tailed) = <0.0001.

